# Functionalized bead assay to measure 3-dimensional traction forces during T-cell activation

**DOI:** 10.1101/2020.09.23.310144

**Authors:** Morteza Aramesh, Simon Mergenthal, Marcel Issler, Birgit Plochberger, Florian Weber, Xiao-Hua Qin, Robert Liska, Georg N. Duda, Johannes B. Huppa, Jonas Ries, Gerhard J. Schütz, Enrico Klotzsch

## Abstract

When T-cells probe their environment for antigens, the bond between the T-cell receptor (TCR) and the peptide-loaded major histocompatibility complex (MHC) is put under tension, thereby influencing the antigen discrimination process. Yet, the quantification of such forces in the context of T-cell signaling is technically challenging. Common approaches such as traction force microscopy (TFM) employ a global readout of the force fields, e.g. by measuring the displacements of hydrogel-embedded marker beads. Recent data, however, indicated that T-cells exert tensile forces locally via TCR-enriched microvilli while scanning the surface of antigen-presenting cells. Here, we developed a traction force microscopy platform, which allows for quantifying the pulls exerted via T-cell microvilli, in both tangential and normal directions, during T-cell activation. For this, we immobilized specific T-cell activating antibodies directly on the marker beads used to read out the hydrogel deformation. Microvilli targeted the functionalized beads, as confirmed by superresolution microscopy of the local actin organization. Moreover, we found that cellular components, such as actin, TCR and CD45 reorganize upon interaction with the beads, such that actin forms a vortex-like ring structure around the beads and TCR is enriched at the bead surface, whereas, CD45 is excluded from bead-microvilli contacts.

**Significance statement:** During the antigen recognition process, T-cells explore and probe their environment via microvilli, which exert local pushes and pulls at the surface of the antigen presenting cell. It is currently believed that these forces influence or even enable the antigen recognition process. Here, we describe the development of a platform, which allows us to quantify the magnitude and direction of traction forces exerted locally by T cell microvilli. Simultaneous Ca^2+^ imaging was used to link the measured forces to the overall T cell activation status. Superresolution microscopy resolved the contact sites of bead-microvilli contact at the nanoscale: cells contacted beads via actin vortex-like structures, which excluded the phosphatase CD45 from the contacts.

## Introduction

T-cell receptors (TCRs) on the surface of T-cells recognize antigenic peptides on major histocompatibility complexes (pMHCs) of antigen presenting cells (APCs) with exquisite sensitivity and specificity (1–3). During the recognition process, the cellular interface – termed immunological synapse – undergoes substantial dynamic reorganizations, with T-cell microvilli contacting and searching the APC’s surface for cognate antigen (4–6). It is still unclear, however, how the molecular binding events between pMHC and the TCR are transduced into intracellular signals (7). Mechanical force has recently been identified as an important parameter in the antigen recognition process (8, 9): On one hand, tensile forces affect the interaction kinetics between TCR and pMHC (10–13), which appears to accentuate differences between agonistic and non-agonistic peptides (14, 15); On the other hand, there are indications that the TCR acts as a mechanosensor, which converts tensile forces into conformational changes, thereby affecting intracellular signaling (16, 17). In line with this, T-cells were shown to push and pull against model APCs (18–20). *Vice versa*, application of force via optically trapped pMHC-functionalized beads triggered T-cell activation, preferentially when forces were applied tangentially to the T-cell surface (21).

Different methods have been applied to quantify tensile forces directly in the synapse between a T-cell and an activating surface. DNA-based molecular tension sensors indicated the presence of pulling forces larger than 12 pN, when TCRs were engaged with antigenic glass-immobilized pMHCs (21, 22). Our own single molecule measurements using peptide-based force sensors revealed slightly lower forces around 5 pN (23). When measured collectively via traction force microscopy, TCR-exerted sheer forces of up to 150 pN were observed (8, 24). While these measurements allowed for quantification of tensile forces down to the level of individual bonds, they provide only indirect access to the directionality of these forces; particularly, forces acting perpendicularly to the T-cell surface have remained unknown. In addition, previous methods did not account for the highly localized force application by T-cells, which involves pulling on the APC surface via microvilli (4).

Here, we report the development of a traction force microscopy platform for the 3-dimensional quantification of tensile forces acting during T-cell activation. Compared to existing methods the platform shows two key advantages. Firstly, we generated well-defined nanoscopic points of force application via functionalized beads. The size of the beads of 200 nm diameter was chosen to match the width of microvilli (25). We used beads coated with anti-CD3 as specific TCR-ligand; in addition, we measured the influence of local co-stimulation via anti-CD28, which was expected to enhance signaling (9). Secondly, the beads were coupled to the surface of elastic gels, offering the possibility for 3-dimensional readout of their displacements. The platform allows for force measurements with a resolution of 56 pN, 76 pN and 93 pN for tangential, normal and total forces, respectively. In addition, the T cell activation status was inferred from simultaneous Ca^2+^ measurements. Using this platform, we were able to directly quantify forces exerted via single T cell microvilli during the activation process.

## Results and discussion

### Functionalized bead assay for T-cell activation

We selected a cytocompatible and photopolymerizable hydrogel system based on Polyethylene (glycol) diacrylate (PEGDA, 10 kDa), which has been widely used for regular cell culture and *in situ* cell photo-encapsulation studies (26). Compared to traditional polyacrylamide gels, PEGDA gels represent significantly lower cytotoxicity, comparable reactivity and minimal protein adsorption (27). The bead assay for activation of T-cells is shown schematically in **Figure 1a**. We designed it with respect to two main aspects:

i. The 3-dimensional displacements of the beads should be directly convertible into normal and tangential forces by solving the inverse stress-strain relationship (28). For this it is critical that individual beads can be treated as independent objects, which can be achieved by choosing sufficiently low bead densities so that their mutual distances are larger than the diameter of the contact area between bead and hydrogel surface (29, 30). In our experiments, the number of beads interacting with a single T-cell was in the range of 30 to 90 and the average attachment area was 175 μm^2^, yielding average nearest neighbor distances of 0.7 - 1.2 μm (31). For the chosen bead diameter of 200 nm beads can hence be treated as independent objects for traction force characterizations (32). We used neutravidin-conjugated beads, which were bound to the surface of photo-crosslinked, biotinylated poly(ethylene glycol) diacrylate (PEGDA) gels.
ii. Forces exerted by T-cells onto the surface should act strictly locally via the functionalized beads. We coupled biotinylated anti-CD3 and/or anti-CD28 to the beads to trigger the TCR’s ε-subunit alone or in combination with the co-stimulatory protein CD28. By choosing beads of 200 nm diameter we aimed at matching the size of naturally occurring microvilli, with which T-cells contact the surface of APCs during the search for antigen (5, 6, 33). Using neutravidin-coated beads makes the assay flexible, as beads could be functionalized also with different biotinylated antibodies or peptides to stimulate the target cells.

**Figure 1:**
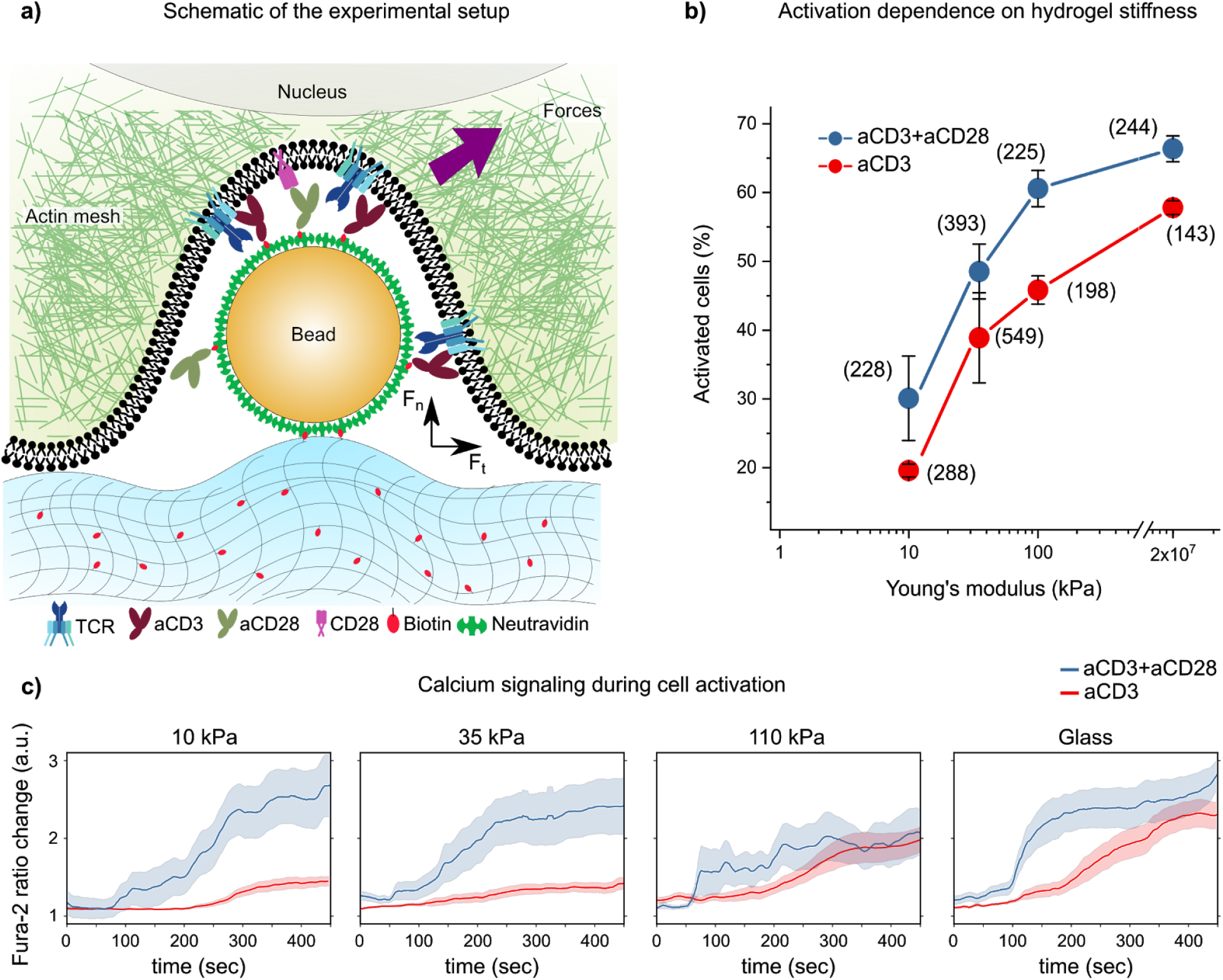
Setup for T-cell activation and Ca^2+^ imaging. **(a)** Schematic of the traction force microscopy assay. Fluorescent neutravidin-coated beads with a diameter of 200 nm were coupled to UV-crosslinked biotinylated PEGDA hydrogels. Beads were functionalized with anti-CD3 and anti-CD28 prior to seeding the cells. **(b)** Effect of hydrogel stiffness on T cell activation. The percentage of activated T cells, defined by a minimal Fura-2 ratio of 1.5, was recorded for PEGDA hydrogels of different stiffness. Data obtained at 20 GPa stiffness correspond to experiments, in which beads were coupled directly to the glass surface. Red and blue data points correspond to experiments performed with beads coated with anti-CD3 only, or with a mixture of anti-CD3 and anti-CD28 antibodies. The number of analyzed cells is given in brackets. **(c)** Kinetics of Ca^2+^ signaling. Fold change of Fura-2 ratio is plotted over time for the different PEGDA hydrogel stiffnesses used in (b), for anti-CD3 with (blue) or without (red) anti-CD28. Fura-2 ratio at the time of the cell first touching down till the onset of Ca^2+^ release (slope increasing) was averaged to calculate Fura-2 ratio change over time. Solid line shows the average over a minimum of 10 cells, shaded areas the standard error of the mean (s.e.m.).

In this paper we showcase the performance of our platform using Jurkat T cells as a model system; early T-cell activation was assessed via the Ca^2+^-sensitive ratiometric dye Fura-2 (34). We first validated the functionality of the platform by quantifying T-cell activation efficiency on gels of different stiffness (**Figure 1b**). Consistent with previous literature (35), we observed a pronounced increase in the fraction of activated cells with increasing stiffness of the PEGDA gel; eventually, beads immobilized directly on poly-d-lysine-coated rigid glass surfaces yielded maximum activation efficiency (data points at 20 GPa stiffness for glass (36)). The delay between surface contact and T cell activation was longer for soft gels compared to stiff surfaces (**Figure 1c**), with the presence of CD28 further reducing this delay. Also this finding was expected, as it is known that stronger stimuli yield faster Ca^2+^ responses (37).

### Combined 3-dimensional Traction Force Microscopy and Calcium imaging

To investigate more closely the generation of 3-dimensional forces in the context of early T-cell signaling, we used substrates of medium stiffness (35 kPa). Beads were imaged via fluorescence microscopy, and their displacements were determined with a precision of 5 nm in *xy* and 30 nm in *z*-direction. Bead displacements were translated into forces as described in the Methods section. Without cells, the determined bead positions showed only small variations, corresponding to force fluctuations of 56 pN, 76 pN and 93 pN for tangential, normal and total forces, respectively (**Figure 2d, control**). We used these residual fluctuations to define the force resolution of the platform.

**Figure 2:**
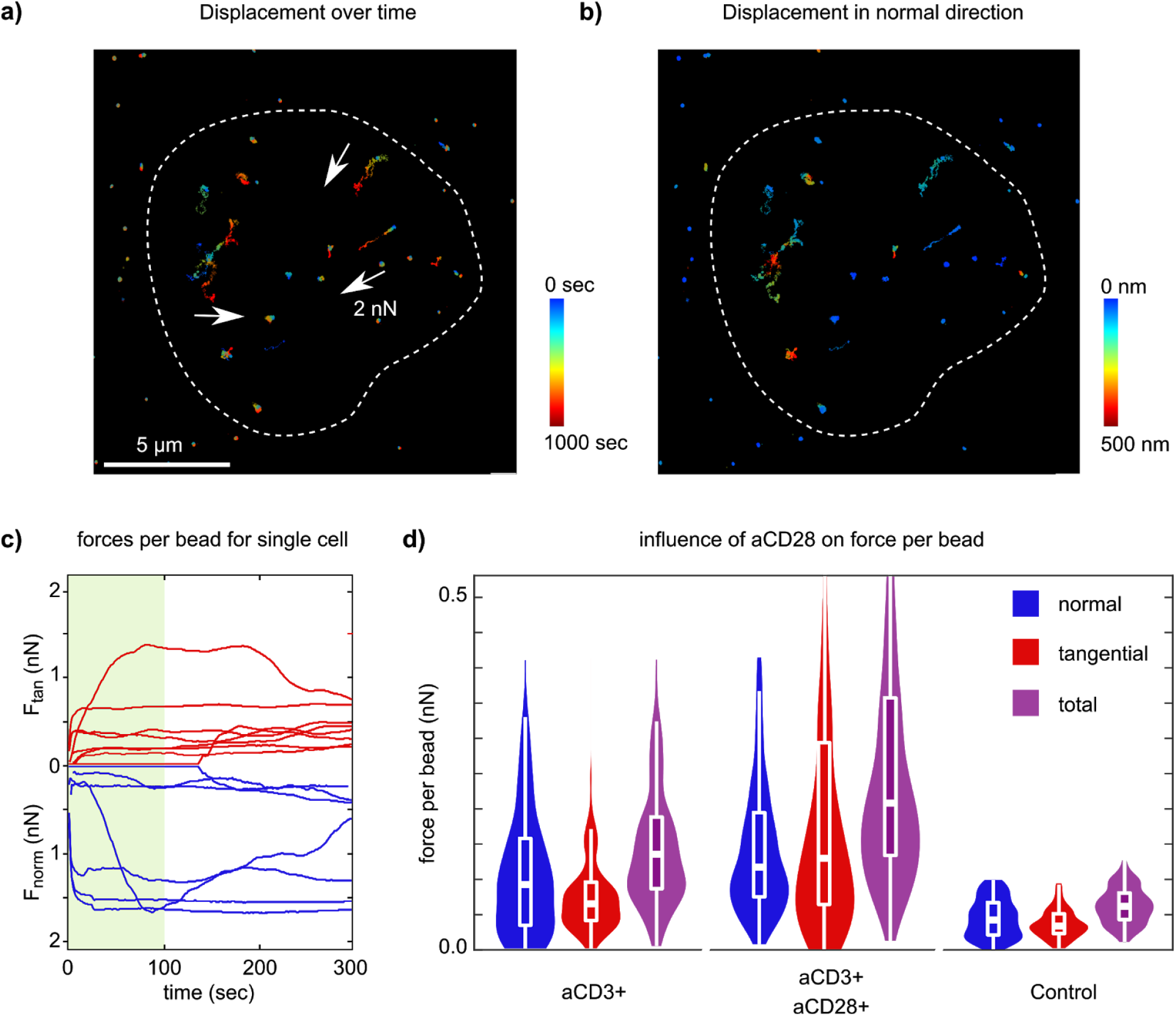
Traction force measurements. **(a)** Bead displacements are plotted over time for an exemplary cell. We used a PEGDA hydrogel with a Young’s modulus of 35 kPa and beads coated both with anti-CD3 and anti-CD28. The left panel shows tangential bead displacements, with color-code indicating time. The white dashed line shown in (**a, b**) indicates the cell outline, derived from the Fura-2 signal. The white arrows represent a bead displacement equivalent to 2 nN. **(b)** Same cell as shown in (**a**), with color code indicating the z-displacements of the beads. **(c)** Normal (blue) and tangential (red) forces experienced by beads underneath the cell are shown within the first 300 s upon cell attachment. Data correspond to the same cell as in panel **(a, b)**. Green shading indicates the time prior to Ca^2+^ flux, defined by a 1.5-fold increase in the Fura-2 ratio. **(d)** The Violin/Whisker-Box plots summarize all recorded forces at the time point of Ca^2+^ flux, with blue, red and purple indicating normal, tangential, and total forces. As control, we determined the maximal force, beads without cell contact were exposed to, representing thermal fluctuations and precision of the bead assay. We defined here as force resolution the mean plus standard deviation, yielding 56 pN, 73 pN and 93 pN for tangential (red), normal (blue) and total (violet) force, respectively. Data represent at least ten cells and 160 beads per condition.

In a typical experiment, T-cells were seeded onto the platform, and we measured both 3D bead displacements and Ca^2+^ traces simultaneously over several minutes (**Figure 2**). Exemplary time traces of the force evolution per bead during the process of T-cell activation are shown in **Figure 2c**; single beads experienced forces up to 500 pN. For a global analysis, we determined the forces acting on individual beads at the time-point of T-cell activation, which we defined by the maximal Ca^2+^ flux for the cell analyzed (**Figure 2d**). Interestingly, tangential forces showed no correlation with normal forces, indicating that the two directions are largely uncoupled (**Figure S1**). T-cells exerted substantially higher forces to anti-CD3/anti-CD28 co-functionalized beads than to beads functionalized only with anti-CD3 (**Figure 2d**), in line with the observation of Bashour *et al. (8)*.

Finally, since T-cells can exert tensile forces exclusively via the beads, our assay allows us to calculate the accumulated force exerted by single T-cells. Exemplary data are shown in **Figure S2**, where we plotted the accumulated force versus the Ca^2+^ response as parametric representation. Not surprisingly, increased force generation correlated with increased Ca^2+^; when Ca^2+^ levels declined at later stages, also forces were reduced again. In total, T cells exerted up 10 nN of force onto the stimulating beads. Given a pulling force of 5-10 pN exerted by single TCR molecules (22, 23), we estimate between 1000 and 2000 TCR molecules involved in the activation process.

### Nanoscale actin ring formation at T-cell-bead contacts

We next aimed at studying the mechanisms of force generation during T-cell activation in more detail. For this, we imaged filamentous actin via expression of tractin-cyOFP (38) (**Figure 3**). We employed structured illumination microscopy (SIM) to visualize the actin distribution at the cells’ basal membrane next to the beads at sufficiently high resolution. In order to perform SIM, however, the PEGDA hydrogel had to be replaced by a material with high refractive index (39); we selected the PDMS-based QGel920 featuring a stiffness of 18 kPa, similar to PEGDA of medium stiffness.

**Figure 3:**
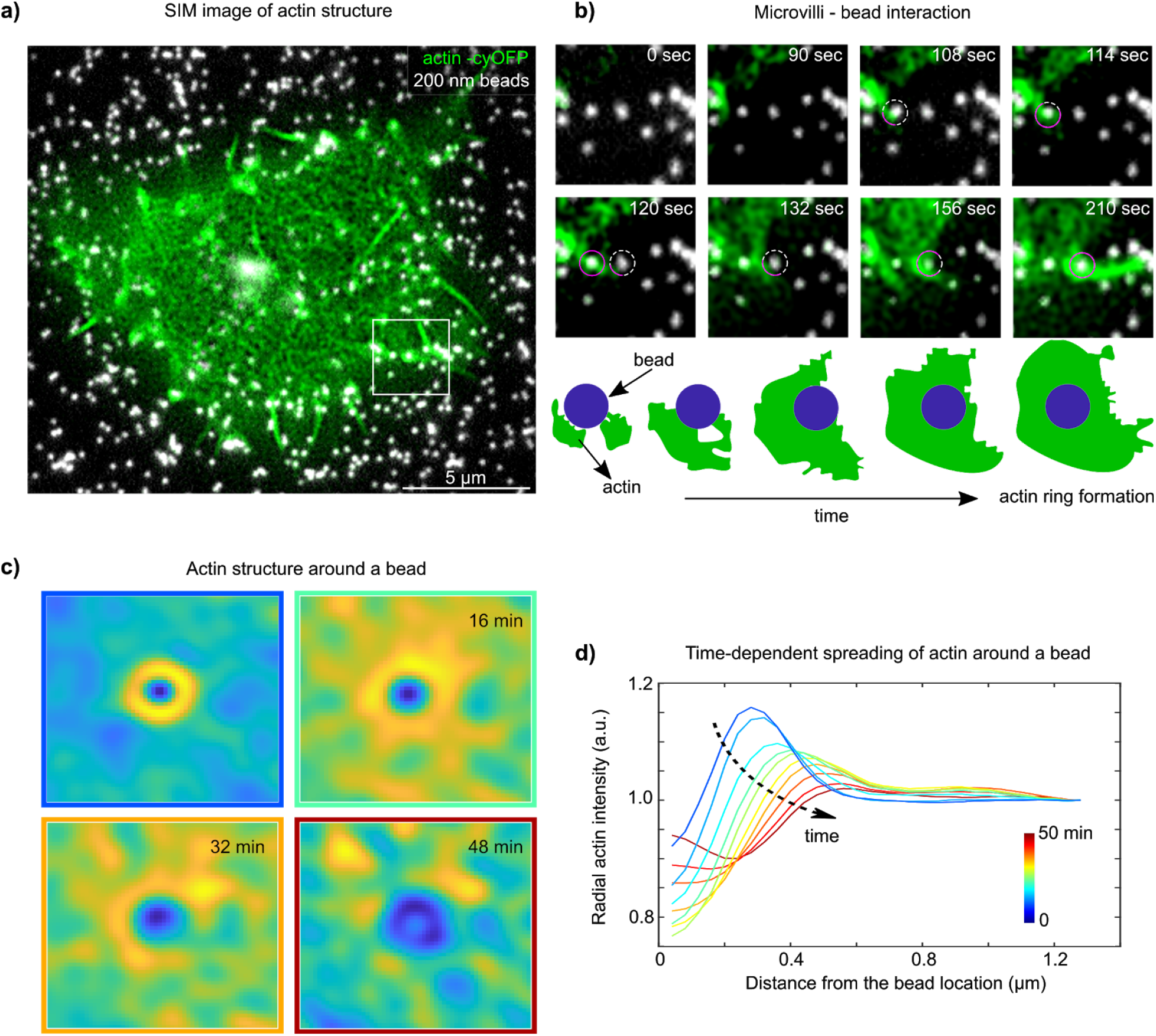
Actin ring formation. **(a)** SIM image of cellular actin (green) overlaid with the calculated bead positions (white). The experiment was performed with anti-CD3/anti-CD28 coated beads. **(b)** The time course of contact formation between a single microvillus and a single bead is exemplarily shown for the detail indicated by the white box in **(a)**. At time point 108s the microvillus-bead contact is established. Within 6s, the microvillus has wrapped around the bead. Over the next two minutes an additional bead was contacted by a microvillus. The schematics below depict the temporal evolution of actin enrichment around the second bead that was contacted by the microvillus. **(c)** The averaged actin intensity around the beads is shown over time. The n = 76 beads were included in the analysis. **(c)** The radial actin intensity with respect to bead positions is plotted as time evolution. Additional data of 10 cells fixed and stained with phalloidin A647 after 15 min, 30 min and 60 min with similar behavior are shown in **Figure S3.** The movie representing the data can be found in **Movie S1**.

SIM images indicate the formation of actin rings around the anti-CD3/anti-CD28 functionalized beads (**Figure 3a**). For a global analysis of all structures, we segmented the actin images into regions around individual beads and averaged over all beads within the cells’ attachment area (**Figure 3c**). Directly after cell attachment an actin ring is formed around the beads (**Figure 3b, left**), reminiscent of actin vortices described previously for cortical actin (40). Closer inspection of the ring morphogenesis, however, revealed surprising details: typically, single microvilli contacted a single bead tangentially, which – within a few tens of seconds – initiated the engulfment of the bead (**Figure 3b**). The distance from the beads’ center to the actin ring increased over time, and eventually the actin ring completely disappeared at late stages (**Figure 3c, Movie S1**).

To quantify the images, we plotted the radial distribution over time (**Figure 3d** and **Figure S3**). To avoid artifacts arising from overlapping signals, beads with a mutual distance smaller than 200 nm were excluded from the analysis. Immediately after T-cell spreading the actin ring had a radius of ~275 nm and a width of ~250 nm (**blue curve**), further supporting the interpretation that single microvilli were wrapping around individual beads. During this process, the interaction between anti-CD3 on the bead surface and the TCR the microvillus surface stabilizes a strongly bent conformation of the actin bundle in the microvillus core. Given the flexural stiffness of actin rods of ~50 pN/nm (41), single microvilli can easily account for the observed tangential forces of a few 100 pN acting on single beads. At later stages, we observed a shift to 500nm radius and a much broader and less dense actin distribution (**red curve**), indicating a massive rearrangement of the membrane morphology close to the bead. Together, the morphogenesis, size, and the stabilization by actin (5) confirm the formation of microvilli-like protrusions, which were contacting single beads.

### TCR bead enrichment and CD45 exclusion on bead-microvilli contacts

Finally, we were interested whether molecular rearrangements of signaling proteins accompany the observed formation of microvilli-bead contacts. Particularly, we were interested in the distribution of the large phosphatase CD45 relative to the bead surface: previously it was hypothesized that size-mediated CD45 exclusion from TCR-ligand contacts shifts the ITAM phosphorylation-dephosphorylation balance, thereby triggering TCR signaling (6).

Cells were fixed at different times of activation and labeled with respective antibodies for TCR and CD45 **(Figure 4a)**. We applied Airyscan microscopy (42) in order to resolve the molecular organization of the TCR and CD45 as well as the positions of anti-CD3/anti-CD28 coated beads.

**Figure 4.**
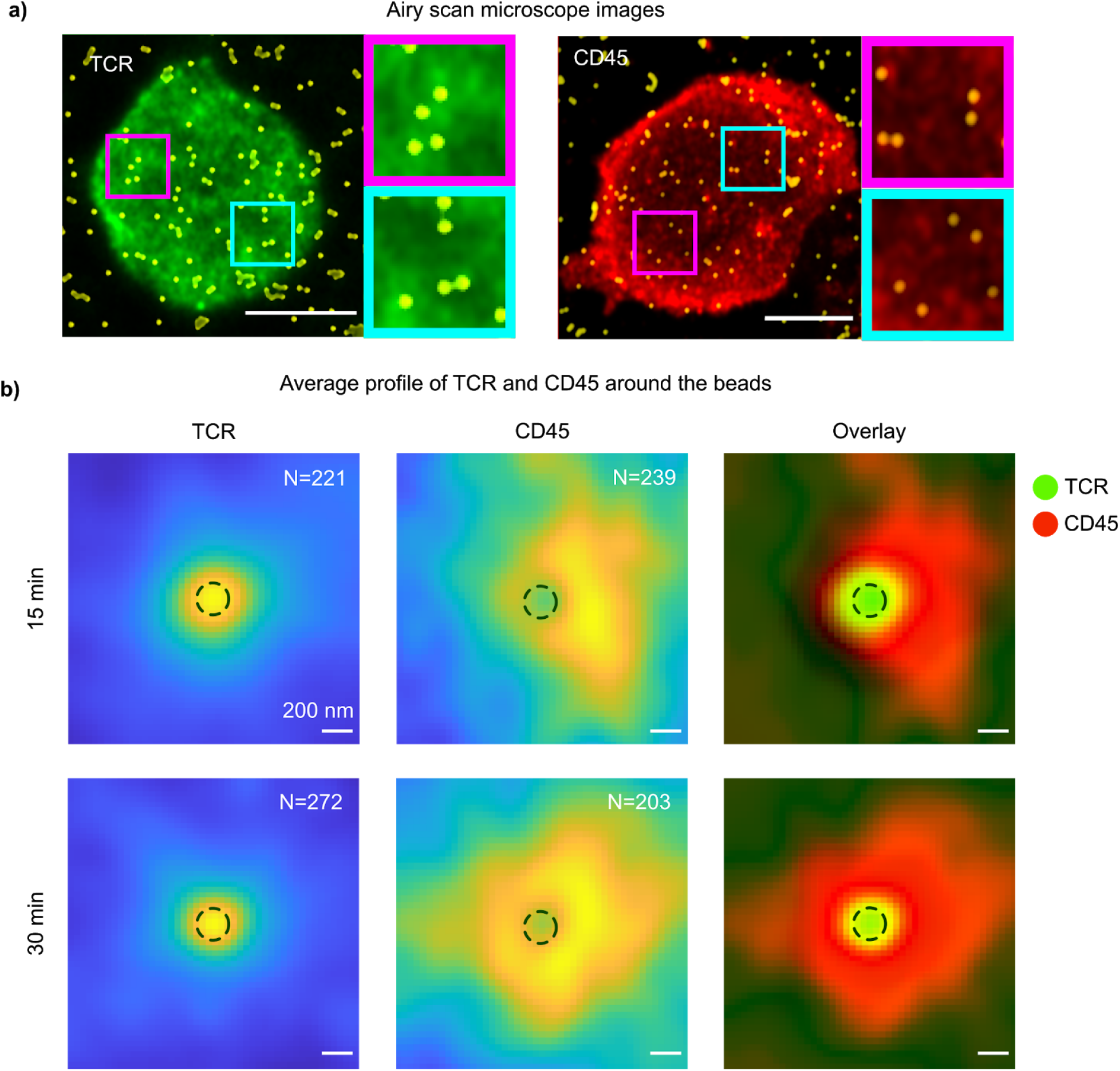
Molecular organization of TCR and CD45 with respect to the beads. Data were obtained using Airy scan microscopy on samples fixed at 15 min or 30 min after T-cell seeding. Experiments were performed using anti-CD3/anti-CD28 loaded beads. In **(a)** beads in yellow are overlaid with the TCR (left, green) and CD45 (right, red) signal; the image corresponds to fixation 15 min post seeding. Two magnified regions are shown on the right side. Scale bar is 5 μm. **(b)** Average intensity profile of registered beads for the TCR- (left) and the CD45-channel (middle) for 15 min (top row) and 30 min (bottom row) (N indicates the number of analyzed beads, with at least 5 cells analyzed). Right column shows the overlay. Scale bar is 200 nm. Dotted line is shown as a guide for the eye to indicate the region of the activating bead.

For quantitative analysis, we segmented the images in **Figure 4a** and determined the protein distribution with respect to the beads’ positions. To avoid artifacts arising from overlapping signals, beads with a mutual distance smaller than 200 nm were excluded from the analysis. In **Figure 4b** we show the TCR- and CD45-distribution obtained from at least 200 registered beads. As expected, we observed the TCR to be associated with the bead surface, irrespective of the time point of fixation. In contrast, CD45 was efficiently excluded from the beads; the effect became more pronounced at later stages of T cell activation.

## Conclusion

In summary, by combining classic with pointillistic TFM, we present a versatile assay to measure and analyze cellular forces in the context of T-cell activation, offering a high potential for functional imaging by modification of bead functionalization. We showed that Ca^2+^ release as one early T-cell activation marker correlates with the mechanical stiffness of the underlying substrate, suggesting the suitability of the method for studying interfaces of cells with novel materials. By implementing fluorescence microscopy, we established 3D traction force microscopy with a 5 nm lateral and 30 nm vertical resolution, enabling high resolution mapping of tangential, normal and total force down to 56 pN, 76 pN, and 93 pN, respectively. Superresolution microscopy revealed initial microvilli-bead contacts, which transformed within seconds to more complex engulfment of the beads, including the formation of vortex-like actin structures. Since the biotinylated ligands can be easily swapped, this methodology and bead assay is applicable also to the study of different cellular signaling mechanisms involving surface-anchored ligands, as in cell adhesion or intercellular contacts.

## Materials and Methods

### Surface Modification and Preparation

Glass coverslips (CS-18R15, Warner Instruments) were either surface functionalized using plasma cleaning for activation followed by vapor phase silanization with Dichlorodimethylsilane (Sigma-Aldrich), resulting in a hydrophobic surface for sandwiching the curing hydrogel or with 3-(Trimethoxysilyl)propylmethacrylate (Sigma Aldrich) resulting in free methacrylate groups for crosslinking of PEGDA hydrogels to the surface.

### PEG

Polyethyleneglycol (PEG, M_n_~10 kDa) was purchased from Sigma-Aldrich (Austria). A water-soluble photoinitiator, Irgacure 2959 (I2959) purchased from BASF (Germany). A heterofunctional PEG-based linker (Biotin-PEG5k-Thiol, Biotin-SH) was purchased from Nanocs Inc (USA). All other reagents were purchased from Sigma-Aldrich and used as received unless otherwise noted.

### PEG Acrylation

PEG diacrylates (PEGDA, 10 kDa) were synthesized according to a generic protocol described by Hubbell et al. (43). Briefly, PEG (10 g, 2 mmol -OH) was dried by azeotropic distillation in 150 mL of toluene for 2 h. After cooling to 40 °C under argon, triethylamine (1.4 mL, 5 equivalents) was added. Afterwards, acryloyl chloride (0.7 mL, 4.5 eq.) was added drop wise. The reaction was stirred overnight at room temperature under argon. The resulting solution was filtered over the alumina bed. The filtrate was subsequently neutralized with sodium bicarbonate and then concentrated by rotary evaporation. The crude product was re-dissolved in 20 mL of dichloromethane and precipitated in cold diethyl ether. After drying in vacuum, PEGDA was obtained as white solid at ~85 % yield. Acrylation degree of PEGDA was x approx. 95 %, as confirmed by ^1^H-NMR analysis **(SI, Figure S4)**. ^1^H-NMR (CDCl3): δ (ppm): 6.40 (2H, m, CH_2_=CH-), 6.20 (2H, m, CH_2_=CH *trans*), 5.80 (2H, m, CH_2_=CH *cis*), 3.80-3.30 (280 × 4 H, s, -CH_2_-CH_2_-). It has to be noted that the efficiency of radical-mediated thiol-acrylate conjugation is as high as 95%, (44) which was known to be higher than that of thiol-acrylamide conjugation (45).

### Photo-Rheology

A plate-to-plate time-resolved photo-rheometer (Anton-Paar MCR-301, **SI, Figure S5)** was used to monitor the light-induced gel formation and quantify the bulk mechanical properties of PEGDA hydrogels. Filtered UV-VIS light (320–500 nm, Omnicure S2000) was directed from the bottom of the rheometer through a glass plate to the sample. Light intensity at the cure place was 10 mW cm^−2^ as determined by an Ocean Optics USB 2000+ spectrometer. After a 60 s blank period, light was triggered to irradiate the samples. Real-time measurements were made in oscillation mode, at 25 °C, 10% strain, 10 Hz and 20 μm gap thickness. Strain and frequency sweeps were performed before and post the polymerization to verify the linear response regime. Plateau value of the storage modulus (G’) of the photopolymerized gels was termed as the bulk gel stiffness.

### Hydrogel Preparation

Briefly, biotin-presenting PEGDA hydrogel layers with a thickness of 20 μm were prepared using a multistep procedure. First, PEGDA and Biotin-SH were dissolved in PBS solution of 0.5% Irgacure (I2959), achieving a final concentration of 30% and 0.16 %, respectively. The resultant solution corresponds to PEGDA gels with elastic moduli of ~100 kPa. In order to access lower stiffness, the 30 % precursor solution was further diluted with 0.5 I2959/PBS solution to a lower PEGDA content of 15 % (for 35 kPa) and 5 % (for 10 kPa), respectively. Second, 30 μL of the precursor solutions were transferred to the central area of a cover slip (diameter: 30 mm), which was pre-functionalized with methacrylate groups for covalent fixation (see above). The obtained specimen was then mounted on the top of the glass plate of a photo-rheometer setting **(SI, Figure S6**). The rheometer platform was lifted down to keep the sample thickness of 20 μm. Finally, the samples were exposed to UV irradiation for 300 s while at the same time performing photo-rheology measurements in order to measure bulk stiffness of the hydrogel.

For SIM and AiryScan microscopy we used QGels920 components A and B (CHT, Quantum silicones) with refractive index of 1.51, in a ratio of 1:5 arriving at a stiffness of 18 kPa using AFM tip indentation **(Figure S7)** (39). The surface was further modified using UV irradiation for 3 min, Sulfo-NHS-LC-biotin (ThermoFisher) dissolved to 1 mg/mL in a Phosphate buffer at pH 8.6 and 50 °C Celsius. The biotinylated surface was then incubated with FluoSpheres™ NeutrAvidin™-marked microspheres (200nm in diameter, fluorescent in yellow-green, ThermoFisher, F8774) with 1-5uL in 150uL PBS per hydrogel. Biotinylated CD3 (OKT3, ThermoFisher) and CD28 (CD28.2, BioLegend) specific antibodies were added concomitantly at 5ug/mL, RT and for at least 30 minutes.

### PEGDA hydrogel characterization for TFM measurements

Bulk stiffness measurements accessed by photo-rheometry allow classification of cells’ ensemble behavior, we were interested in the sensation of the local stiffness as the cell senses force on the individual bead level(46). For this, AFM tip indentation (see paragraph below) was used to probe the PEGDA surface stiffness locally and also to test for lateral homogeneity of the hydrogels **(Figure S8, S9)**. For that, BSA-coated AFM tips were scanned over five regions of 90 × 90 μm^2^ with 16 × 16 points of indentation, revealing an average Young’s modulus of 14 ± 5 kPa, 33 ± 14 kPa and 95 ± 19 kPa for the soft, medium and hard hydrogel stiffness maps using a tip velocity of 2 μm/s with a maximal indenting force of 30-400 pN **(see Figure S8)**. Furthermore, in order to test for gel elasticity, we checked for the independence of Young’s modulus from the loading rate of indentations at the same position **(see Figure S9)**, with a slight trend towards material softening at higher speeds between 25 and 500 pN/s. This allowed us to exclude non-elastic stress relaxation contributions to T-cell behavior as was measured to be important for cell size regulation by Mooney et al.(47).

### AFM stiffness measurements

PEGDA and PDMS hydrogels were characterized with Atomic Force Microscopy (NanoWizard 3, JPK, Berlin). Soft silicon nitride AFM probes with nominal spring constants of k ~ 0.01 N/m and 0.02 N/m (MSNL, Veeco Instruments, Plainview, NY) were used for the indentation experiments. The exact spring constant of each cantilever was determined from the thermal noise spectrum (48–50). An example of a force-indentation curve recorded from the hydrogel is shown in (**Figure S10**). Elasticity fits (Hertz models) with conical tip geometry, corrected sample-tip distance and a Poisson ratio of 0.5. were performed using JPK software.

### Superresolution microscopy for bead position combined with Fura-2 Calcium imaging

To obtain bead locations in 3D, we used TIRF microscopy. Measurements were performed using a custom-built single-molecule microscope. A 488-nm laser (200 mW; Coherent) was coupled through a single-mode fiber (QiOptics) into a Polytrope and Yanus (TILL Photonics) mounted on an inverted Zeiss Axiovert 200 microscope. The beam was then focused onto the back-focal plane of a high numerical aperture objective (α–Plan-Apochromat 100×/1.46; Zeiss) for highly inclined illumination (illumination intensity ~ 1 kW/cm2) or TIRF microscopy. Emission light was filtered using appropriate filter sets for Alexa 488. The emission light was imaged with a back-illuminated iXon DU 897 EMCCD camera (Andor Technology Ltd.), which was water-cooled to −80 °C. For functional calcium imaging the excitation was switched between 488nm and 340nm/380nm white light Halogen lamp, using the Polytrope (Till Photonics). This allowed us to subsequently measure calcium and bead position every 500 ms.

We estimated the lateral drift to be smaller than 50 nm/h, while drift correction was implemented at the level of the analysis software (see below). We observed no change in the widths of individual beads not exposed to cells during experiments, hence z-drift was not corrected.

### Localization Analysis

The data were analyzed as previously described (51). Briefly, smoothing, non-maximum suppression, and thresholding revealed the possible locations of fluorescent beads. Selected regions of interest were fitted by a pixelated Gaussian function and a homogeneous photon background with a maximum likelihood estimator for Poisson distributed data using a freely available fast GPU (graphics processing unit) fitting routine on a GeForce GTX1080 Ti (Nvidia).

We typically acquired 2000 frames for the reconstruction and data analysis of bead position. Lateral drift was corrected based on bead positions. The displacements corresponding to each time point were averaged using a robust estimator that was interpolated by a spline and used to correct the position of each localization. We estimated that the residual errors for the corrected positions were about 5 nm.

The localization data were rendered using the Thomson blurring. Briefly, each localization is represented as a 2D Gaussian function with a width according to the precision of the respective localization determined from the fitted number of photons and background(52). All analysis software was written in MATLAB, based on the Single Molecule Analysis Platform (SMAP) by Ries lab (53). All quantitative analyses were done using the list of localization coordinates and not the processed images; the latter are only presented for visualization.

### Bead displacement to force calculations

To derive forces in normal and tangential direction we used the following formulas for spring coefficients (30)

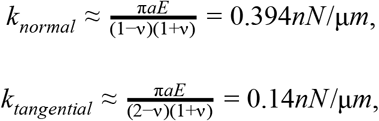

where we estimate the Poisson ratio *ν* with 0.45 for our PEGDA hydrogel(27), the bead radius of a = 100 nm and a Young’s modulus E of 35 kPa for the medium stiffness hydrogel in **Figure 2**. Forces in normal and tangential direction were calculated with:

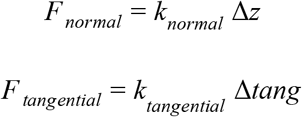

where 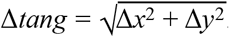. Total forces were calculated from forces in normal and tangential direction.

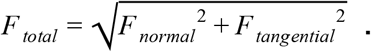

### Single Cell Calcium Imaging Analysis

All live-cell and fixed cell experiments were done with Jurkat T-cells (E6-1, ATCC), live-cell experiments were performed at 37 °C adjusted by an objective heating system (OKO lab). Cells were loaded with 10 μM Fura-2/AM (Molecular Probes®) in complete RPMI 1640 at room temperature for 60 min, washed with HBSS (Invitrogen) and flushed onto stimulatory surfaces on the stage of the above described custom built microscope. Cells were illuminated using a halogen lamp at 340/10nm and 380/10nm (polytrope, Till Photonics) and fluorescence emissions collected via a α–Plan-Apochromat 100×/1.46; Zeiss. Movies were acquired at a frame rate of 0.5 frames per second, the emission ratio of 340/380 nm was determined after background correction and preprocessed in Jupyter notebooks. Individual cells were selected manually and Ca2+ time traces were calculated after background correction. For noise reduction and elimination of outliers, a median filter with a window size of 10 time points was applied. The background for ratio of Fura-2 increase over time was calculated by averaging Fura-2 intensity from moment of cell touching down until Fura-2 increase.

### Immunostaining

Jurkat cells were seeded on the hydrogel surface (50,000 cells per cm^2^) and incubated at 37 °C and 5% CO_2_ for 15-60 min. The cells were then fixed in pre-warmed 4% PFA (Formaldehyde, Polysciences, Inc) and 0.1% Triton-X in Cytoskeletal buffer for 7 min. The Cytoskeletal buffer consists of10 mM MES pH 6.1, 150 mM NaCl, 5 mM EGTA, 5 mM glucose and 5 mM MgCl_2_. For actin staining, the cells were pre-fixed with 0.5% PFA in Cytoskeletal buffer for 1 min. The samples were then washed with PBS three times and incubated with 0.01% NH_4_Cl in PBS for 10 min to reduce the autofluorescence. The samples were then washed with PBS for three times and were blocked with 2% BSA in PBS for 1 h at room temperature before staining. Phalloidin AlexaFluo 647 from Thermo Fisher Scientific in 1:200 dilution was used for actin staining. Anti-CD45 antibody (F10-89-4, Abcam and EP322Y, Abcam) and anti-CD3 antibody (clone SP7, ab16669, Abcam) in 1:200 dilutions were used to stain CD45 and TCR, respectively. The samples were then incubated with AlexaFluor 647-conjugated secondary antibodies for 1 hour at room temperature in the dark. The samples were then washed with PBS three times before mounting with Prolong gold antifade reagent (Molecular Probes).

### Airyscan Microscopy

For fixed cells imaging Zeiss LSM 880 Airyscan microscope was used to acquire images using 63x 1.4NA Oil DIC Plan-Apochromat objective. Argon (488 nm) and HeNe (633 nm) lasers were used to image the beads and the cells, respectively. Zen 2012 software was used as the interface for data acquisition.

### Structured Illumination Microscopy

For super-resolution images, structured illumination microscopy (Zeiss Elyra) was used. Diode lasers with 488 and 561 nm were used for illumination, with a cyOFP specific 570 - 650 nm filter, where fluorescent beads FluoSpheres™ NeutrAvidin™-marked microspheres (200nm in diameter, fluorescent in yellow-green, ThermoFisher, F8774) were measured using a 495 - 550 nm emission filter. The signals were recorded by the Andor iXON 3 885 camera. We used an alpha Plan-Apochromat 100× with 1.46 NA objective. The data was acquired with Zeiss Software, and was reconstructed and drift corrected with custom written Matlab code. We estimated the resolution of the microscope to be 160 nm in lateral and 350 nm axial direction for the actin structure, while bead position was estimated *via* wide field imaging and localization as described above. The cells were transfected with cyOFPtractin for 24h prior cells seeding on hydrogels. Hydrogels were temperature equilibrated with culture media (RPMI) at 37 °C Celsius for at least 1 hour prior cell seeding.

## Acknowledgement

We thank Wes Legant and Michael L. Smith for general discussions and helpful advice regarding TFM and Daniel Nieves for advice with surface functionalization. We thank MZ Lin for kindly providing the cyOFP tractin plasmid. EK thanks for financial support from Human Frontiers Science Program RGY0065/2017, FEBS Long term fellowship throughout the onset of the project and ARC DECRA fellowship. MA funding for SNF Spark. GJS and JBH acknowledges funding from the Vienna Science and Technology Fund (WWTF) project LS13-030.

## Supplementary Information

**Figure S1:**
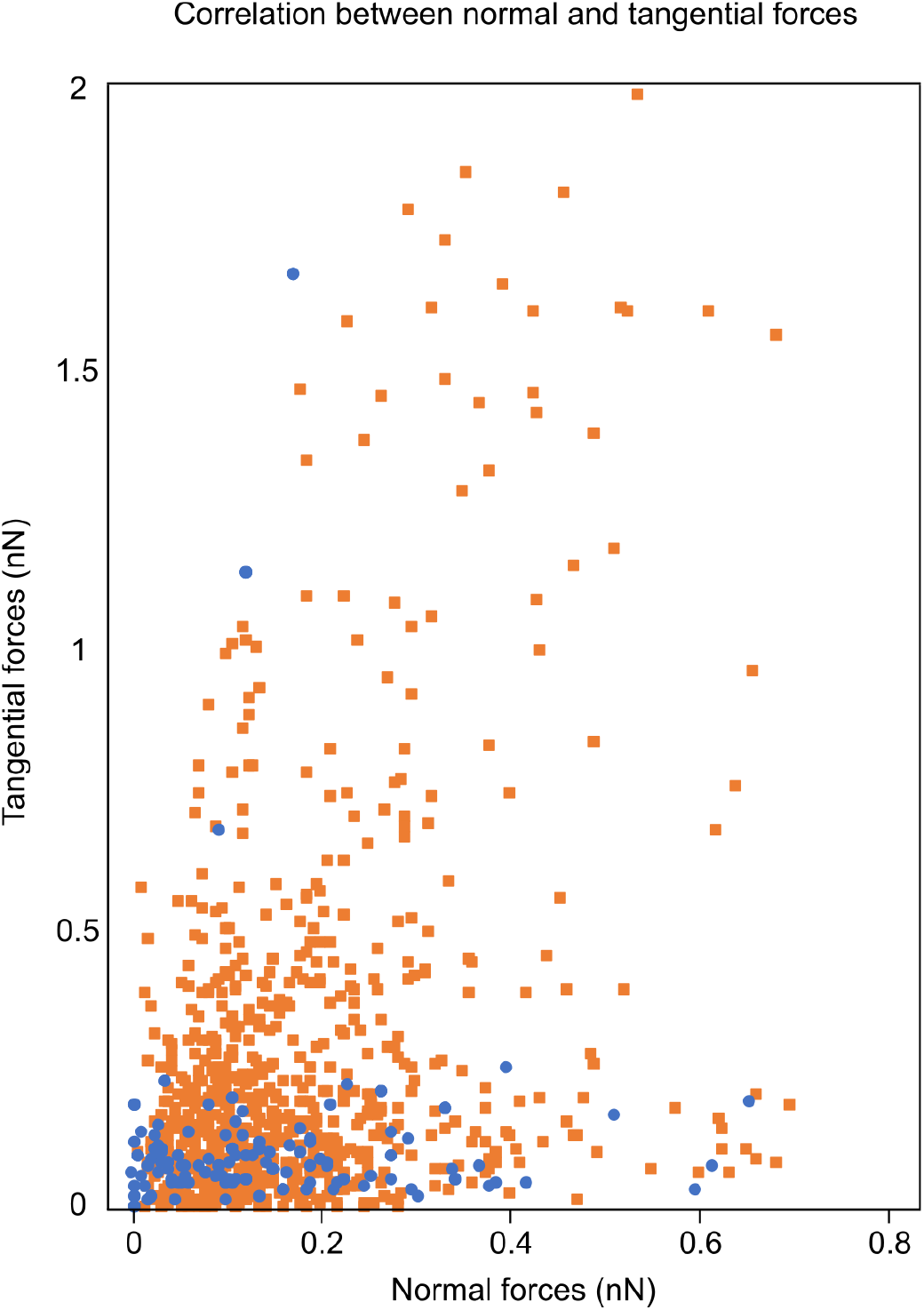
Correlation between normal and tangential forces. Data points from Figure 2d are plotted as scatter plots, yielding a Spearman correlation of 0.09 for bead coated with anti-CD3 only (blue) and 0.25 for beads coated both with anti-CD3 and anti-CD28 (orange).

**Figure S2:**
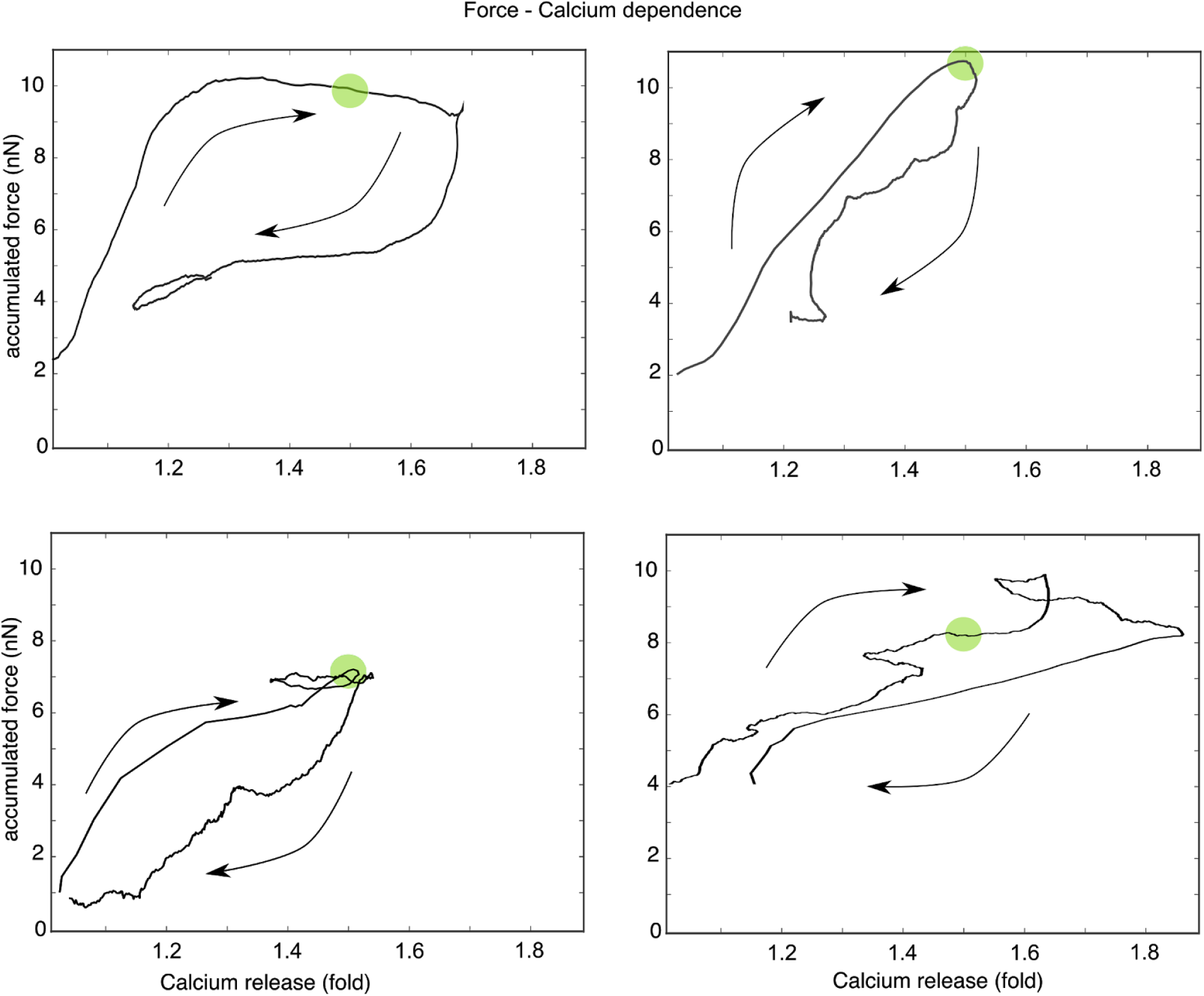
Four exemplary parametric force versus Fura-2 ratio plots are shown. T-cells were exposed to anti-CD3 and anti-CD28 coated beads. The measurements on the upper left corner represent data shown in Figure 2A-C. Green dot represents the time when Ca^2+^ ratio reached 1.5 fold increase.

**Figure S3:**
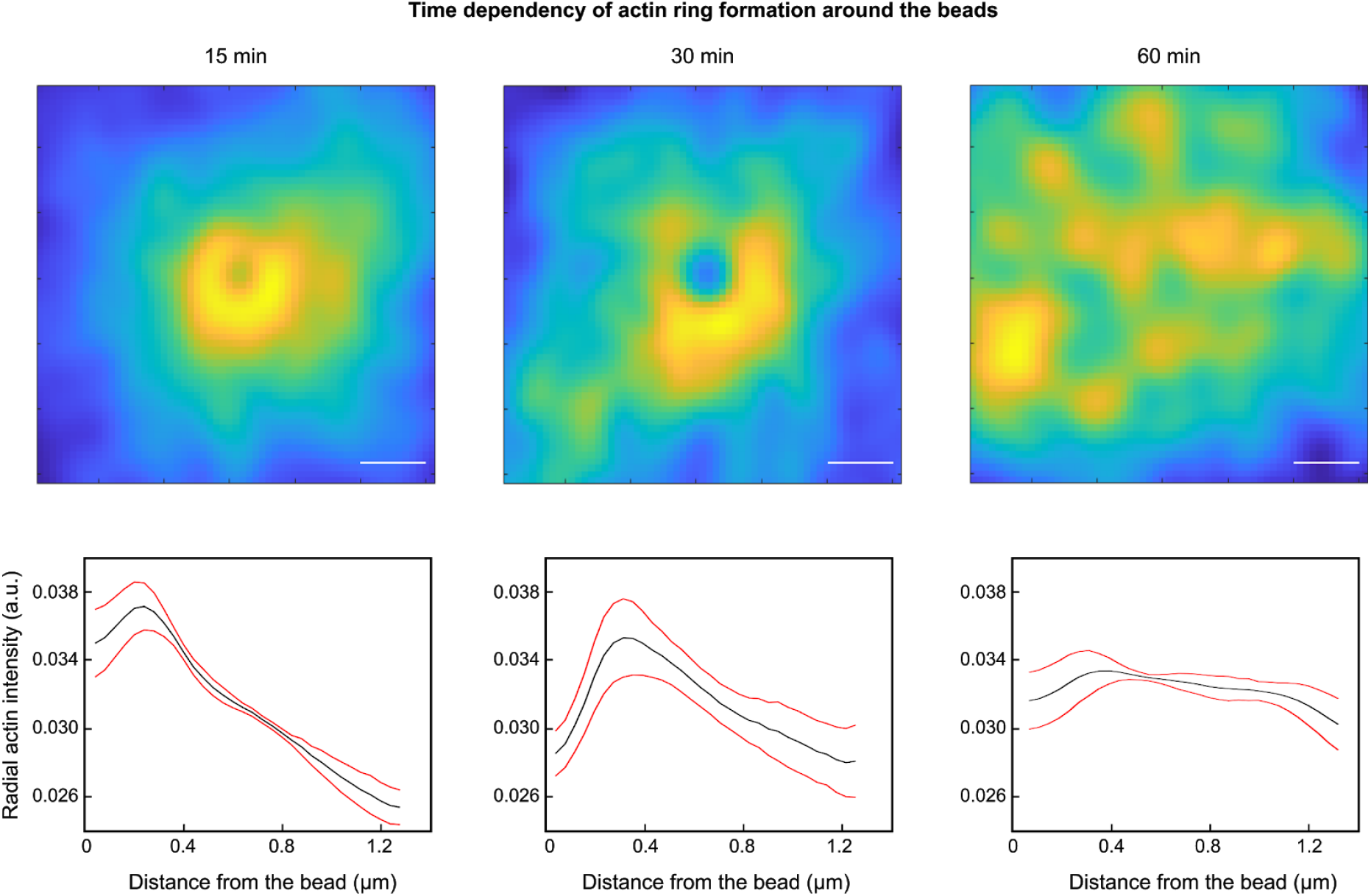
Additional experiments demonstrating actin ring formation. **(a)** The averaged actin intensity around activating beads for at least 10 cells is shown for 15 min, 30 min and 60 min after T-cell activation on anti-CD3/anti-CD28 coated beads. In contrast to Figure 3, cells were fixed before staining with phalloidin-A647. Scale bar is 400 nm. **(b)** The average radial actin intensity is plotted as a black line for the same time points as in (a); red lines indicate with standard deviation derived from ten different cells with at least 30 beads.

**Figure S4.**
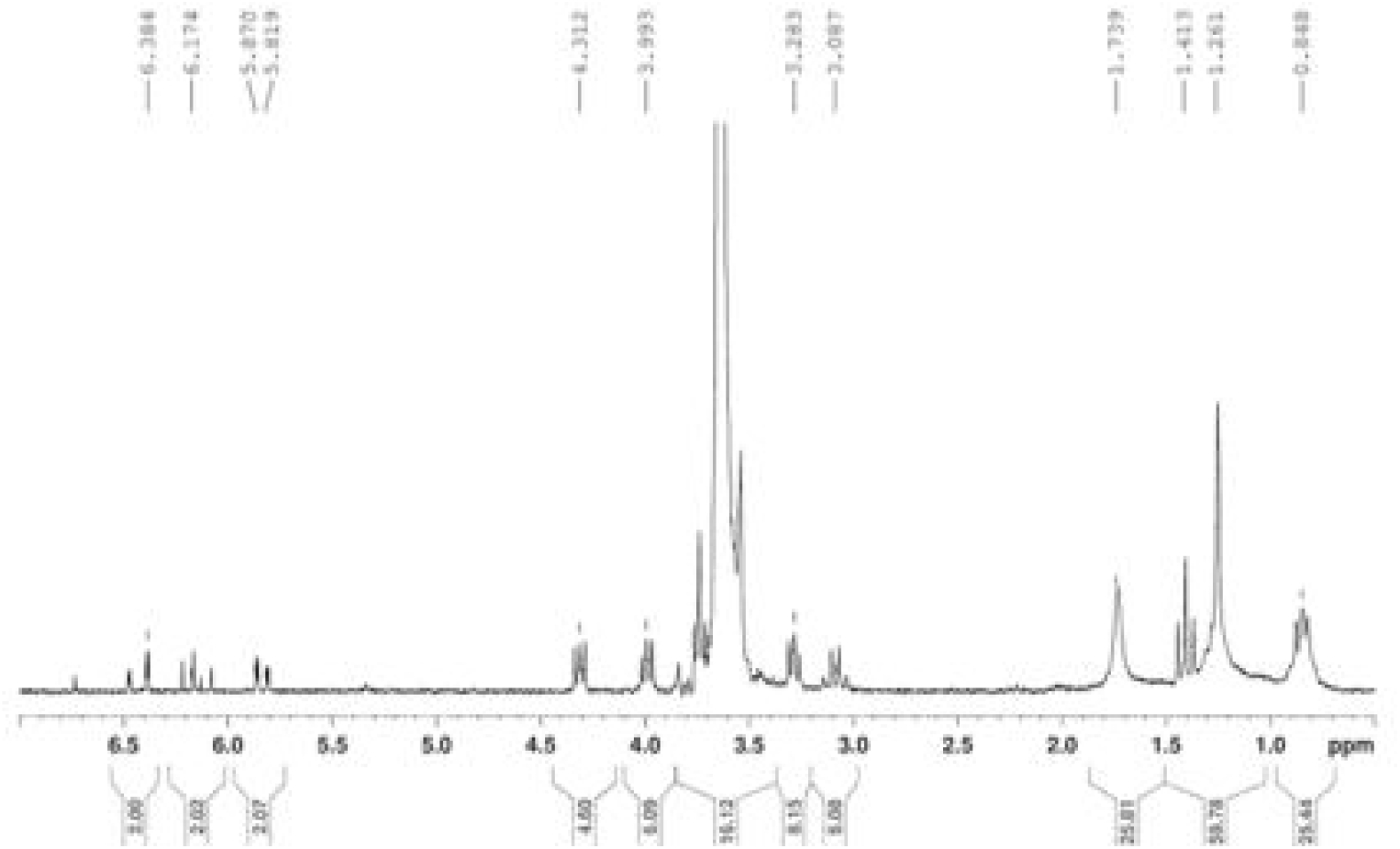
^1^H-NMR spectrum of PEGDA-10k (200 MHz, CDCl_3_).

**Figure S5.**
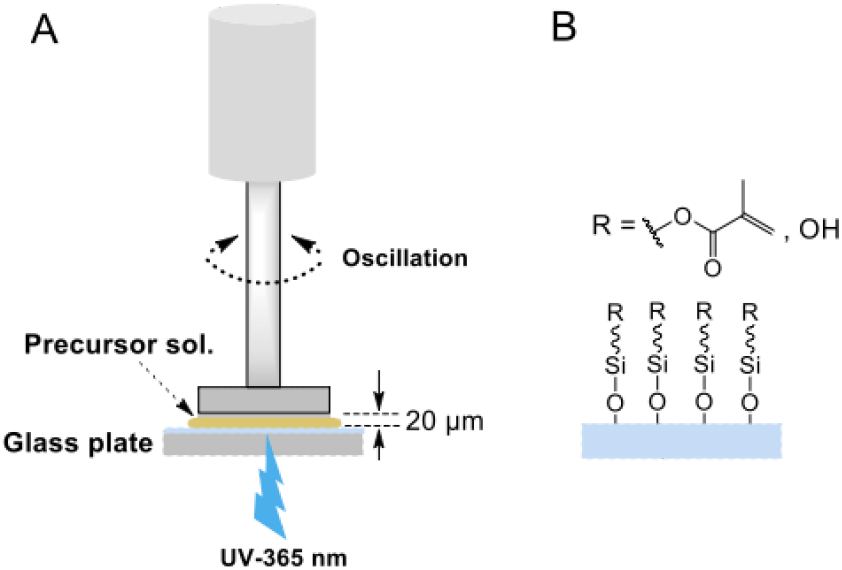
**(A)** Schematic showing the preparation of hydrogel thin layers on a photo-rheometer as well as the *in situ* oscillatory measurement of gel stiffness. **(B)** Schematic of the surface chemistry of methacrylated glass as the substrate for covalent fixation of PEG hydrogels.

**Figure S6:**
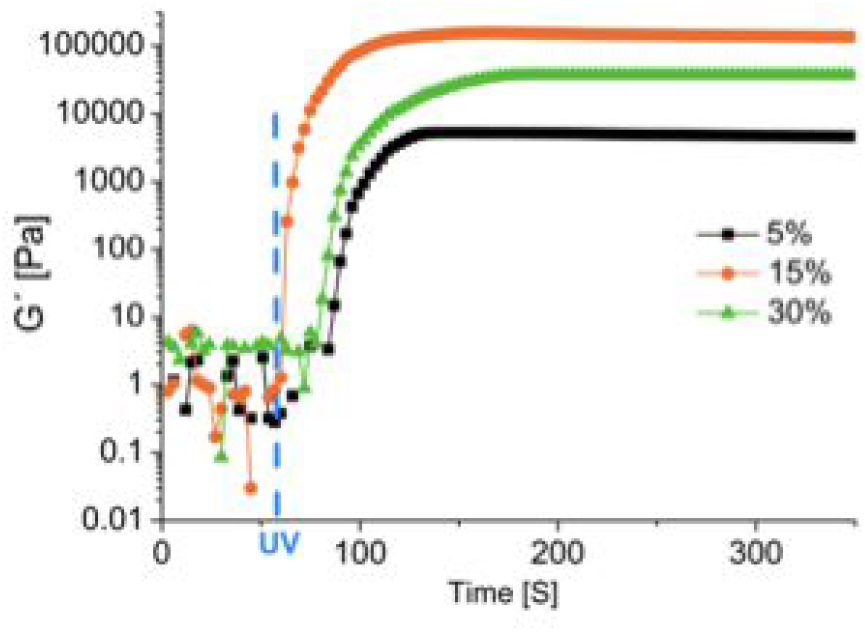
Photo-Rheology measurements for the different stiffness hydrogels. Young’s modulus is plotted over the time of the UV crosslinking process. Saturation is reached after approximately 120 seconds at the target moduli defined by the chosen concentrations.

**Figure S7:**
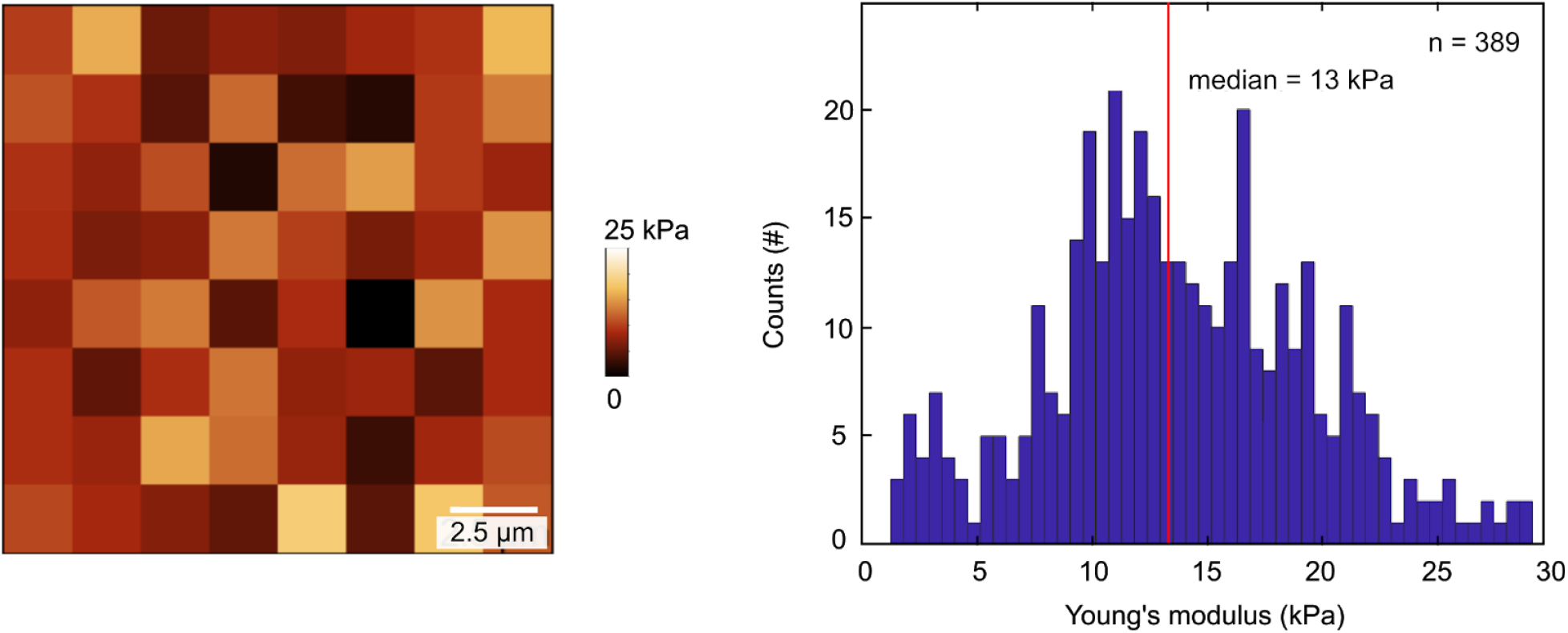
AFM indentation maps to measure Young’s modulus of the Qgel920 based hydrogel. Young’s modulus maps are shown for mixing QGel components A and B (CHT, Quantum silicones) at a ratio of 1:5 arriving at a stiffness of 13.0 ±5.7 kPa for the 20 by 20 μm^2^ area.

**Figure S8:**
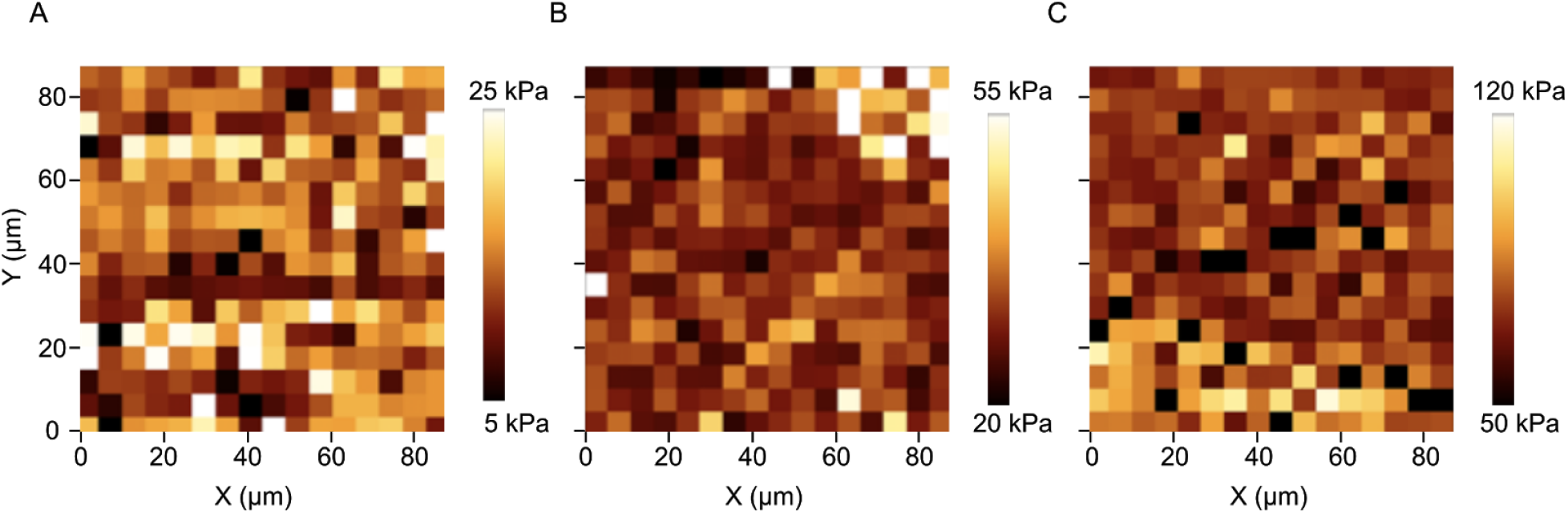
AFM indentation maps to measure Young’s modulus of the hydrogel. Young’s modulus maps are shown for **(A)** soft, **(B)** medium and **(C)** hard PEGDA gels; indentation measurements were performed every 5.6 micrometer. We calculated mean values of 14±5 kPa (**A**), 33±14 kPa (**B**) and 95±19 kPa (**C**).

**Figure S9:**
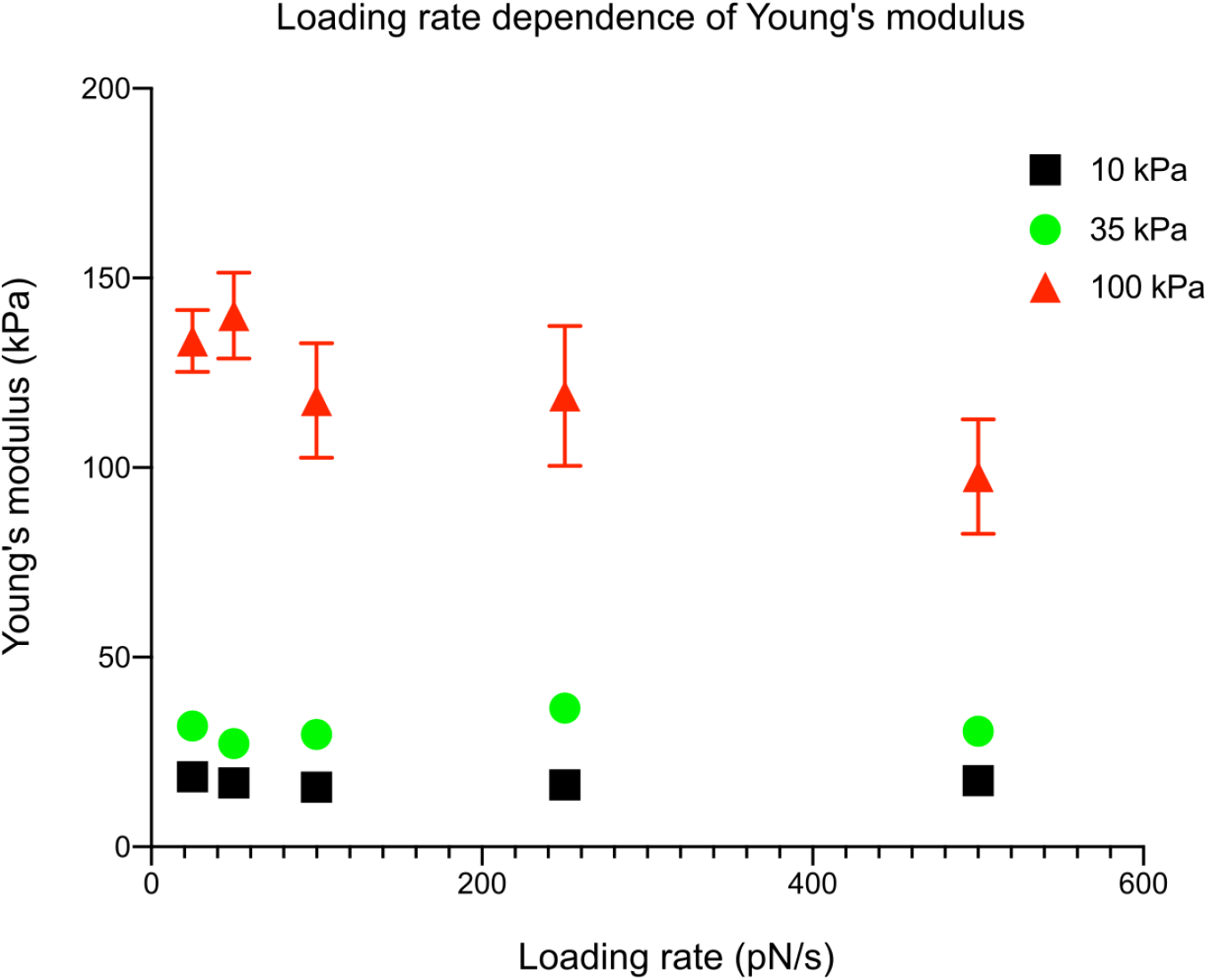
AFM indentation loading rate dependence of different maps to measure Young’s modulus of the hydrogel. Young’s modulus maps are shown for soft, medium and hard PEGDA gels.

**Figure S10:**
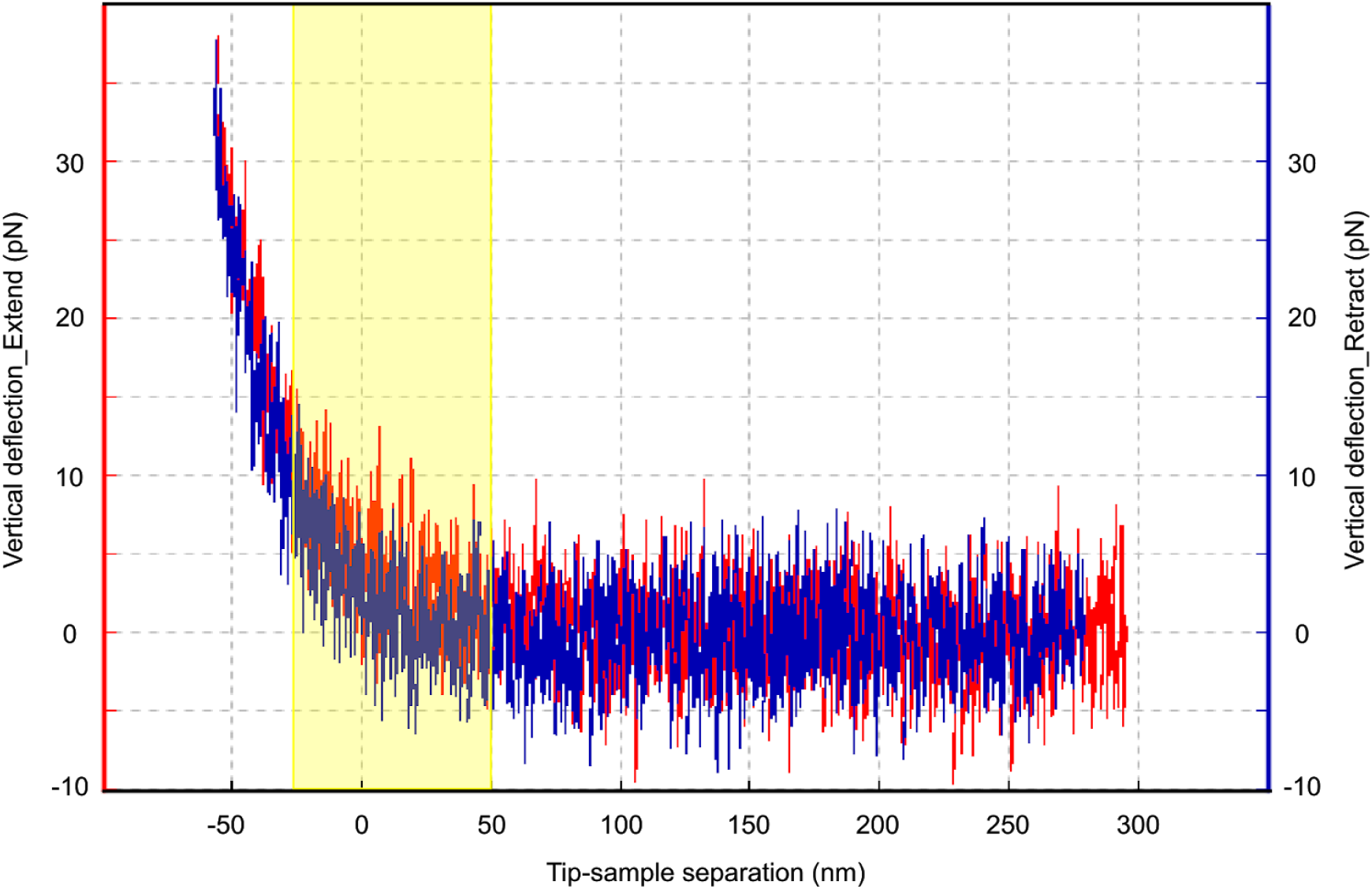
An example AFM tip indentation curve performed on a hydrogel is shown. Approach Curve (Red), Retrace (Blue), Fitted region (Hertz-Model) (Yellow). The measurement was performed using a cantilever with a spring constant of 10 pN/nm, a tip velocity of 1 μm/sec and a set trigger force of 35 pN.

**Movie S1: Actin dynamics during T-cell bead interaction visualized by SIM**. Fluorosphere beads are in grey, actin is visualised with cyOFP-tractin in green. The total length of the movie represents 50 minutes in real time.

